# Hybridization and introgression are prevalent in Southern European *Erysimum* (Brassicaceae) species

**DOI:** 10.1101/2021.11.03.467125

**Authors:** Carolina Osuna-Mascaró, Rafael Rubio de Casas, José M. Gómez, João Loureiro, Silvia Castro, Jacob B. Landis, Robin Hopkins, Francisco Perfectti

## Abstract

**Background and Aims:** Hybridization is a common and important force in plant evolution. One of its outcomes is introgression - the transfer of small genomic regions from one taxon to another by hybridization and repeated backcrossing. This process is believed to be common in glacial refugia, where range expansions and contractions can lead to cycles of sympatry and isolation, creating conditions for extensive hybridization and introgression. Polyploidization is another genome-wide process with a major influence on plant evolution. Both hybridization and polyploidization can have complex effects on plant evolution. However, these effects are often difficult to understand in recently evolved species complexes.

**Methods:** We combined flow cytometry, transcriptomic and genomic analyses, and pollen-tube growth assays to investigate the consequences of polyploidization, hybridization, and introgression on the recent evolution of several *Erysimum* (Brassicaceae) species from the South of the Iberian Peninsula, a well-known glacial refugium. This species complex differentiated in the last 2Myr, and its evolution has been hypothesized to be determined mainly by polyploidization, interspecific hybridization, and introgression.

**Key Results:** Our results support a scenario of widespread hybridization involving both extant and “ghost” taxa. Several taxa studied here, most notably those with purple corollas, are polyploids, likely of allopolyploid origin. Moreover, hybridization in this group might be an ongoing phenomenon, as prezygotic barriers appeared weak in many cases.

**Conclusions:** The evolution of *Erysimum* spp. has been determined by hybridization to a large extent. The adaptive value of such genomic exchanges remains unclear, but our results indicate the importance of hybridization for plant diversification across evolutionary scales.

## INTRODUCTION

Hybridization is widespread across the tree of life, determining the branching and diversification patterns of many taxonomic groups (Rieseberg and Carney, 1998; Coyne and Orr, 2004; Abbott et al., 2013; Arnold, 2016). Because of its pervasiveness, hybridization has been a subject of research for a long time (Stebbins, 1959; Anderson, 1953; Arnold et al., 1999). However, it is only recently, with the advent of next-generation sequencing, that scientists have started to analyze the dynamics of hybridization at the scale of whole genomes, thus rekindling interest in the evolutionary relevance of this phenomenon. Although the patterns of hybridization remain unexplored for many groups, the renewed research efforts have undoubtedly increased our understanding of the role of hybridization in nature (Payseur and Rieseberg, 2016; Goulet et al., 2017; Taylor and Larson, 2019).

Hybridization is particularly relevant for plant evolution, with many plant species showing hybrid origins (Mallet, 2005; Soltis and Soltis, 2009). The evolutionary outcomes of hybridization may vary widely. Interspecific hybridization can hinder speciation and therefore diversification (Mayr, 1992; Schemske, 2000; Mallet, 2005; Saari and Faeth, 2012; Gómez et al. 2015a), but in other cases, hybridization can actually foster the formation of new species (Rieseberg et al., 2003; Stelkens and Seehausen, 2009) or the introgression of novel genetic variation (by hybridization and repeated backcrossing; Anderson and Hubricht, 1938; Anderson, 1953; Rieseberg and Wendel, 1993). In addition, the fusion of genomes between two hybridizing species can lead to changes in ploidy levels (i.e., allopolyploidization; Soltis et al., 2014). There is evidence that introgression might even span ploidy levels (e.g., gene flow between diploid and tetraploid species of Senecio; Chapman and Abbott, 2010), which opens intriguing questions about the interplay of introgression and polyploidization. However, the specifics of how hybridization, introgression, and polyploidization interact to affect the evolution of particular plant groups remain poorly understood. Advancements in genomic sequencing technology and analyses are now making the challenges of characterizing these processes far more feasible, even in recently diverged lineages and taxa.

*Erysimum* L. is one of the largest genera of the Brassicaceae, comprising more than 200 species (Polatschek, 1986), and has been described as a taxonomically complex genus with a reticulated evolutionary history in which polyploidization may have affected the evolution of some clades (Marhold & Lihová, 2006; Turner, 2006; Abdelaziz, 2013; Muñoz-Pajares, 2013). This genus is distributed mainly in Eurasia, with some species in North America and North Africa (Warwick et al., 2006). Notably, more than a hundred species have been described in the Mediterranean region (Greuter et al., 1986) with particular abundance in the Iberian Peninsula, where twenty-one (Polatschek, 1979; Polatschek, 2014) or twenty-three (Nieto-Feliner, 1993; Mateo et al., 1998) species have been described. Most Iberian *Erysimum* species have yellow flowers, but six have purple corollas (Nieto-Feliner, 1993; Gómez et al., 2015b). Interestingly, previous studies suggested that some purple species may have a recent, hybrid, and allopolyploid origin (Nieto-Feliner, 1992, Nieto-Feliner, 1993; Abdelaziz et al., 2014; Gómez et al. 2014). A history of hybridization could further suggest the possibility that the purple flower color has been transferred across the Iberian clade through hybridization and then maintained by natural selection. This scenario would indicate that introgression and polyploidization are intertwined in this group and might have contributed to the adaptive evolution of *Erysimum* spp.

Here we studied signals of hybridization across six species of *Erysimum* (*E. mediohispanicum, E. nevadense, E. fitzii, E. popovii, E. baeticum, E. bastetanum*) that inhabit the Baetic Mountains, an important and dynamic glacial refugium (Médail and Diadema, 2009). The evolution of several plant species has been hypothesized to have been affected by speciation and secondary contacts in this region (Médail and Diadema, 2009; Nieto-Feliner, 2011). The repeated expansion and contraction of ranges and the subsequent cycles of sympatry and isolation might have created conditions for extensive hybridization, introgression, and allopolyploid formation. This species group appears to have differentiated relatively rapidly within the last 2Myr (Osuna-Mascaró et al., 2021). Previous authors have hypothesized that this rapid evolution has been strongly affected by polyploidization and hybridization, as this group spans several ploidy levels, and some species pairs have been reported to produce fertile hybrids (Abdelaziz et al., 2014; Abdelaziz et al., 2021). Species of this group show characteristics that may facilitate ongoing introgression, such as growing in sympatry in some locations and having a generalist pollination system that renders gene flow among different species possible.

The main goal of this study is to disentangle the history of hybridization for the *Erysimum* species complex in the Baetic Mountains. Specifically, we considered both whole-genome effects of hybridization (i.e., the interplay between hybridization and polyploidization) and local, potentially important, introgression of specific genomic regions. Moreover, we also quantified prezygotic barriers among extant taxa to estimate the likelihood of gene flow among them. We test the hypotheses that a) Genomes of this species complex must exhibit signals of multiple hybridization events; b) Some taxa might be allopolyploid, and c) If purple corollas are the product of introgression, hybridization and gene-flow should be detectable, and prezygotic barriers may be weak between (at least some) yellow and purple taxa.

## MATERIAL AND METHODS

### Plant samples

We studied six species in the genus *Erysimum* collected in the Baetic Mountains, South of Spain (Table 1; Figure 1). Specifically, we sampled three different populations for *E. mediohispanicum* (yellow corollas; Em21, Em39, Em71), *E. nevadense* (yellow corollas; En05, En10, En12), *E. popovii* (purple corollas; Ep16, Ep20, Ep27), *E. bastetanum* (purple corollas; Ebt01, Ebt12, Ebt13), and *E. baeticum* (purple corollas; Ebb07, Ebb10, Ebb12), and one population for *E. fitzii* (yellow corollas; Ef01). Some of these species appear in sympatry in some of the sampled localities (e.g., *E. popovii*, Ep20, and *E. mediohispanicum*, Em39; Table 1). Additionally, we sampled one population of *E. lagascae* (Ela07), an allopatric diploid species with purple corollas inhabiting Central Spain, posited as one potential parental species of the Baetic Mountain species studied here (Nieto-Feliner, 1993). We collected fully developed flower buds for transcriptomic analyses (five buds from an individual per population) and leaves for flow cytometry (6-10 individuals per population).

**Table 1.**
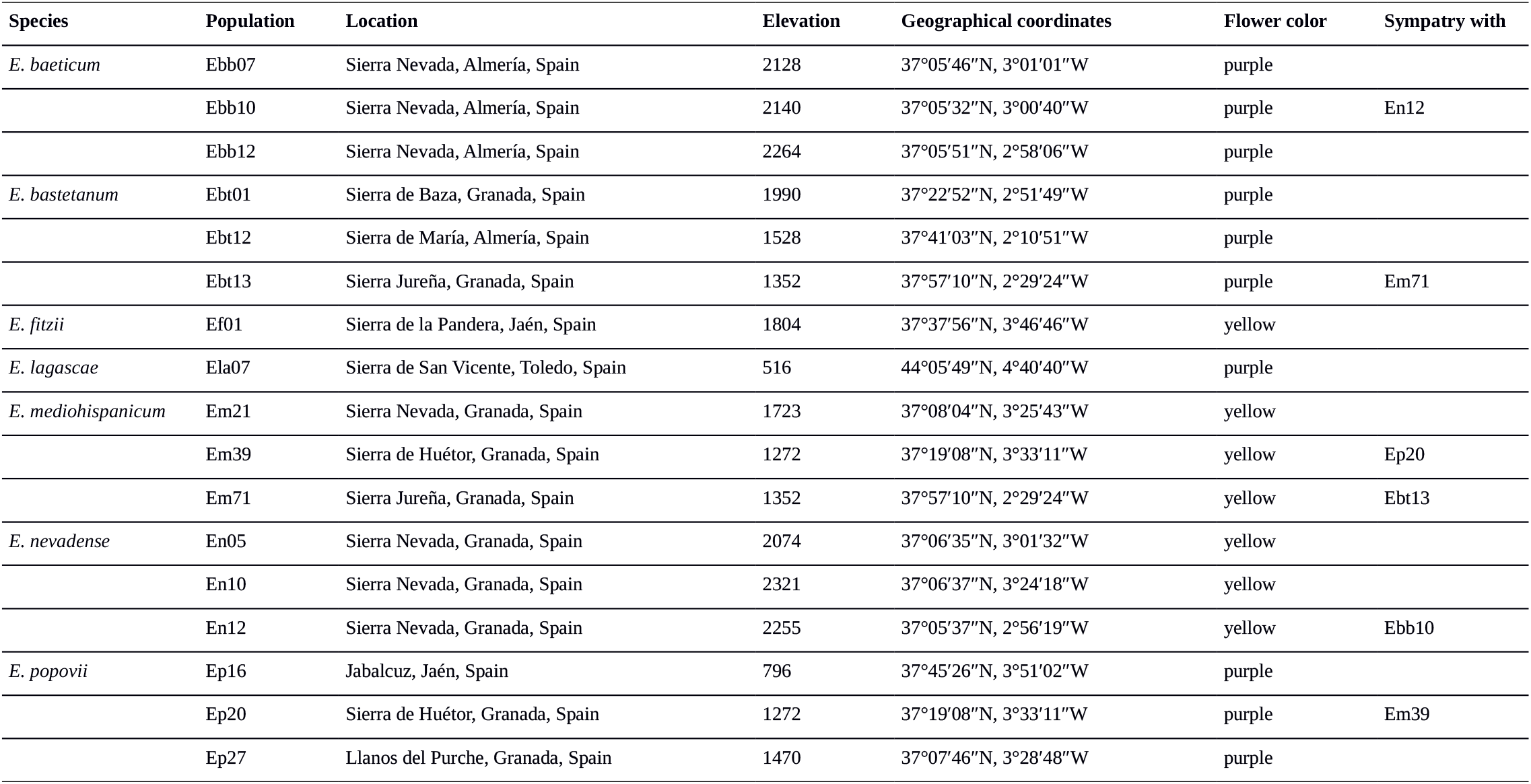
Population code, location, and details of sympatry status for all of the populations sampled.

**Figure 1.**
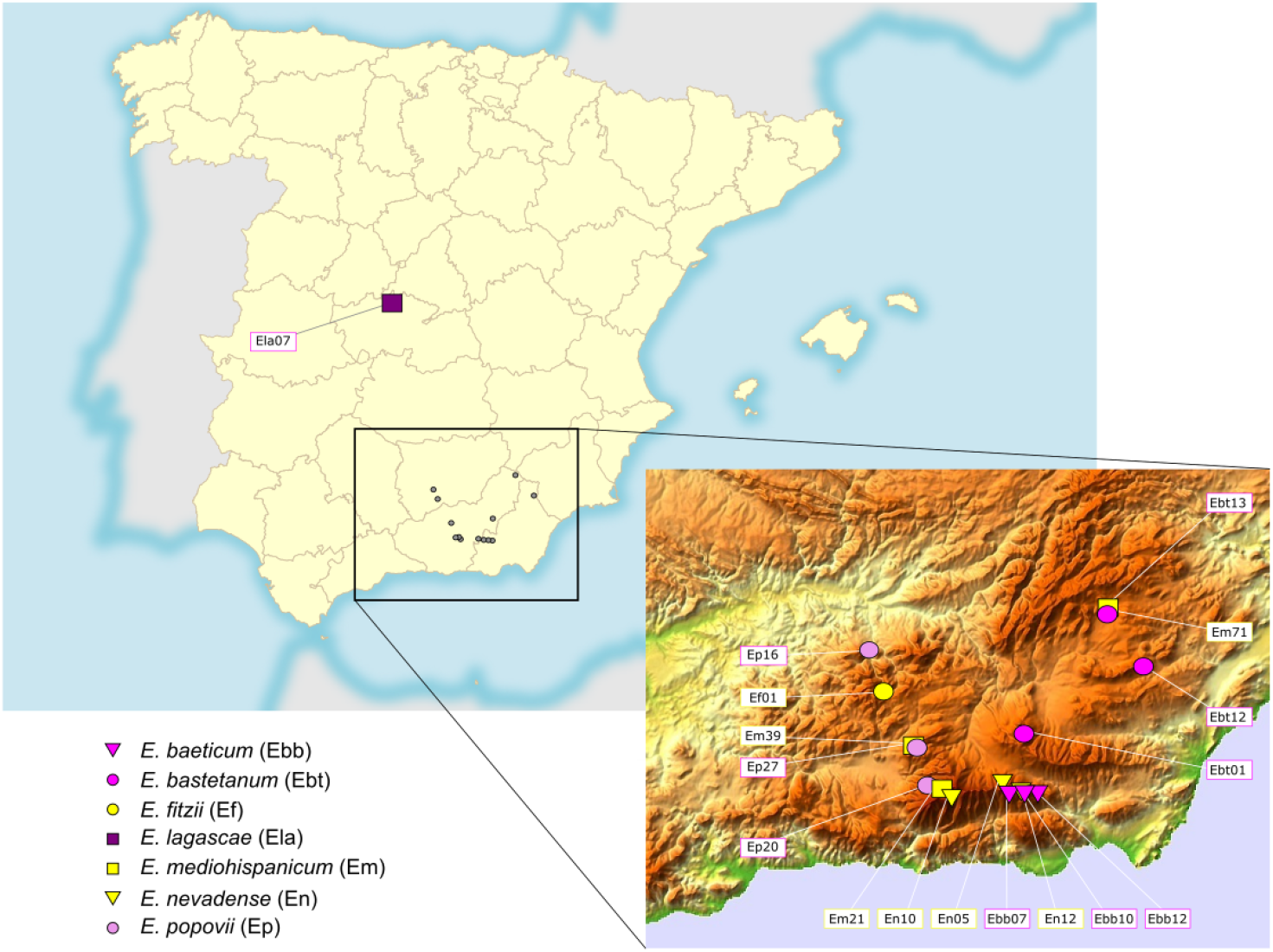
Map of the Iberian Peninsula showing the location of the sampled populations. The insert shows a more detailed map of the Baetic mountains. Purple species are represented with purple symbols and yellow species are represented with yellow symbols. The populations Ebt13 – Em71, Ebb10 – En12, and Em39 – Ep27 represent population pairs located in sympatry.

### Flow cytometry analyses

We used flow cytometry to assess genome size and estimate DNA ploidy levels. Nuclei were isolated from fresh leaf tissues by simultaneously chopping with a razor blade 0.5 cm2 of leaf and 0.5 cm2 of an internal reference standard (Galbraith et al., 1983). We used Solanum lycopersicum L. ‘Stupické’ with 2C = 1.96 pg or Raphanus sativus L. with 2C = 1.11 pg as internal reference standards (Doležel et al., 1992). The nuclei extraction was made on a Petri dish containing 1 ml of WPB buffer (Loureiro et al., 2007). Then, the nuclear suspension was filtered using a 50 µm nylon mesh, and DNA was stained with 50 µg ml-1 of propidium iodide (PI, Fluka, Buchs, Switzerland). Additionally, 50 µg ml-1 of RNAse (Fluka, Buchs, Switzerland) was added to degrade dsRNA. After a 5 min incubation, the samples were analyzed in a Sysmex CyFlow Space flow cytometer (532 nm green solid-state laser, operating at 30 mW). Results were acquired using FloMax software v2.4d (Partec GmbH, Münster, Germany) in the form of four graphics: histogram of fluorescence pulse integral in linear scale (FL); forward light scatter (FS) vs. side light scatter (SS), both in logarithmic (log) scale; FL vs. time; and FL vs. SS in log scale. The FL histogram was gated using a polygonal region defined in the FL vs. SS histogram to avoid debris signals. At least 5,000 particles were analyzed per sample. Only CV values of 2C peak of each sample below 5% were accepted; otherwise, a new sample was prepared and analyzed until quality standards were achieved (Greilhuber et al., 2007). In a few cases, samples produced histograms of poorer quality even after repetition due to the presence of cytosolic compounds. Thus, it was impossible to estimate ploidy level and/or genome size for some samples (Table 2).

**Table 2.** Genome size estimates and DNA ploidy levels obtained in populations of *Erysimum*. The following data are given for each population and ploidy level: mean, the standard deviation of the mean (SD), coefficient of variation (CV, %), minimum (Min) and maximum values (Max) of the holoploid genome size (2C, pg) followed by sample size for genome size estimates (N); DNA ploidy level (2n) and respective sample size (N) for ploidy estimates. DNA ploidy levels: 2x, diploid; 4x, tetraploid; 8x, octoploid; 10x, decaploid. For Ebb12, En05, and En10 samples were not possible to estimate the ploidy levels, and we have used the described in Blanca et al. (1992).

| Species | Population | DNA Ploidy level |  | Genome size (2C, pg) |  |  |  |  |  |
| --- | --- | --- | --- | --- | --- | --- | --- | --- | --- |
|  |  | 2n | N | Mean | SD | CV | Min | Max | N |
| *E. baeticum* | Ebb07 | 8x | 5 | 2.08 | 0.08 | 3.85 | 1.93 | 2.17 | 2 |
|  | Ebb10 | 8x | 6 | 2.07 | 0.09 | 4.35 | 1.93 | 2.17 | 5 |
|  | Ebb12 | 8x | - | - | - | - | - | - | - |
| *E. bastetanum* | Ebt01 | 4x | 4 | 1.06 | 0.06 | 5.66 | 0.97 | 1.10 | 4 |
|  | Ebt12 | 4x | 2 | 1.06 | 0.12 | 11.32 | 0.97 | 1.15 | 2 |
|  | Ebt13 | 8x | 64 | 1.96 | 0.06 | 3.06 | 1.87 | 2.17 | 60 |
| *E. fitzii* | Ef01 | 2x | 3 | 0.44 | 0.004 | 0.91 | 0.44 | 0.45 | 3 |
| *E. lagascae* | Ela07 | 2x | 10 | 0.46 | 0.02 | 4.35 | 0.44 | 0.50 | 10 |
| *E. mediohispanicum* | Em21 | 2x | 2 | 0.44 | 0.01 | 2.27 | 0.43 | 0.44 | 2 |
|  | Em39 | 2x | 21 | 0.46 | 0.02 | 4.35 | 0.43 | 0.49 | 19 |
|  | Em71 | 4x | 59 | 0.98 | 0.04 | 4.08 | 0.93 | 1.13 | 59 |
| *E. nevadense* | En05 | 2x | - | - | - | - | - | - | - |
|  | En10 | 2x | - | - | - | - | - | - | - |
|  | En12 | 2x | 3 | 0.45 | 0.03 | 6.67 | 0.42 | 0.47 | 3 |
| *E. popovii* | Ep16 | 4x | 3 | 0.98 | 0.02 | 2.041.86 | 0.95 | 1.00 | 3 |
|  | Ep20 | 10x | 15 | 2.49 | 0.06 | 2.416 | 2.40 | 2.60 | 10 |
|  | Ep27 | 4x | 39 | 0.96 | 0.04 | 4.17 | 0.92 | 1.05 | 9 |

We obtained the genome size in mass units (2C in pg; sensu Greilhuber et al., 2005) using the formula: sample 2C nuclear DNA content (pg) = (sample G1 peak mean/reference standard G1 peak mean)* genome size of the reference. The ploidy levels were inferred for each sample based on chromosome counts and genome size estimates available for the genus and species.

### RNA extraction and sequencing

Details of the sampling, RNA extraction, and sequencing appear in Osuna-Mascaró et al. (2021). In summary, we stored collected flower buds of each individual in liquid nitrogen until RNA extraction. Floral buds were ground with a mortar and a pestle in liquid nitrogen. We used the Qiagen RNeasy Plant Mini Kit following the manufacturer’s protocol to isolate total RNA from 17 samples (one individual per population; three populations of *E. baeticum, E. bastetanum, E. mediohispanicum, E. nevadense*, and *E. popovii*, and one population of *E. fitzii* and *E. lagascae*). Then, we checked the quality and quantity of the RNA using a NanoDrop 2000 spectrophotometer (Thermo Fisher Scientific, Wilmington, Delaware, United States) and agarose gel electrophoresis. Library preparation and RNA sequencing were conducted at Macrogen Inc. (Seoul, Korea). Before sequencing, the quality of the RNA was analyzed again with the Agilent 2100 Bioanalyzer system (Agilent Technologies Inc., Santa Clara, California, United States), and an rRNA-depletion procedure with Ribo-Zero (Illumina, San Diego, California, United States) was used to enrich mRNA content and to avoid the sequencing of rRNA. Library preparation was performed using the TruSeq Stranded Total RNA LT Sample Preparation Kit (Plant). Sequencing of the 17 libraries (one per individual) was carried out using the Hiseq 3000-4000 sequencing protocol and TruSeq 3000-4000 SBS Kit v 3 reagent, following a paired-end 150 bp strategy on the Illumina HiSeq 4000 platform. A summary of sequencing statistics is shown in Table S1 (Supporting Information).

### Transcriptome assembly and annotation

Details of the read quality control, trimming, and de novo transcriptome assembly and annotation can be found in Osuna-Mascaró et al. (2021). Briefly, we used FastQC v0.11.5 (Andrews, 2010) to analyze the quality of each library’s raw reads. Then, we trimmed the adapters in the raw reads using cutadapt v1.15 (Martin, 2011), and we quality-filtered the reads using Sickle v1.33 (Joshi and Fass, 2011). After trimming, we used FastQC v0.11.5 (Andrews, 2010) again to verify the trimming efficiency. To assemble the resulting high-quality, cleaned reads into contigs, we followed a de novo approach using Trinity v 2.8.4 (Grabherr et al. 2011). Before assembly, each library was normalized in silico to validate and reduce the number of reads using the “insilico_read_normalization.pl” function in Trinity (Haas et al., 2013). Then we used the parameter ‘min_kmer_cov 2’ to eliminate single occurrence k-mers heavily enriched in sequencing errors, following Haas et al. (2013). Candidate open reading frames (ORF) within transcript sequences were predicted and translated using TransDecoder v 5.2.0 (Haas et al., 2013). We performed functional annotation of Trinity transcripts with ORFs using Trinotate v 3.0.1 (Haas, 2015). Sequences were searched against UniProt (UniProt Consortium, 2014), using SwissProt databases (Bairoch and Apweiler, 2000) (with BLASTX and BLASTP searching and an e-value cutoff of 10-5). We also used the Pfam database (Bateman et al., 2004) to annotate protein domains for each predicted protein sequence. Transcripts were filtered through the eggnog (Jensen et al., 2007), GO (Gene Ontology Consortium, 2004), and Kegg (Kanehisa and Goto, 2000) annotation databases.

### Orthology inference

To reduce redundancy, we clustered the translated sequences using cd-hit v 4.6 (Li and Godzik, 2006), following the steps of the pipeline described in Yang and Smith (2014). For the inference of orthologs, we excluded UTRs and non-coding transcripts, using only coding DNA sequences (CDS) in order to avoid the inclusion of sequencing errors (Yang and Smith, 2014). We identified ortholog genes using the OrthoFinder v 2.3.3 pipeline (Emms and Kelly, 2015). In brief, this pipeline first made a BLASTP analysis with the protein sequences as input for searching the orthogroups (a set of potentially orthologs protein-coding genes derived from a single gene in the last common ancestor of all the species sampled), then clustered and aligned the orthologous sequences using MAFFT v 7.450 (Katoh and Standley, 2013) with default parameters. Finally, we obtained the maximum-likelihood phylogenetic gene trees for all orthogroups using IQ-Tree v 1.6.1 (Nguyen et al., 2014). Then, each orthogroup that contained sequenced from all sampled species was used to infer a species tree using STAG v 1.0.0 (Emms and Kelly, 2019). Then, we used DLCpar v 1.1 (Wu et al., 2014) to reconcile the species tree with the gene trees, considering gene duplication, losses, and incomplete lineage sorting (ILS) as potential causes of discordance among trees.

### Phylogenetic reconstruction

We obtained a coalescent species tree using ASTRAL v 5.6.3 (Mirarab et al., 2014) with default parameters. This method reconstructs a species tree from unrooted gene tree topologies. We used the gene trees previously obtained by maximum likelihood by using IQ-Tree v 1.6.1 as input. We used FigTree v 1.4.0 (Rambaut and Drummond, 2012) to visualize and edit the species tree. Then, we compared the alternative tree topologies with the phylogeny obtained from whole chloroplast genome analyses for the same species (presented in Osuna-Mascaró et al., 2021) using the Shimodaira-Hasegawa Test (SH-Test; Shimodaira and Hasegawa, 1999) from the R package phangorn v 2.5.5 (Schliep, 2011). Both phylogenies were also compared visually, plotting them as mirror images with the function cophyloplot, using the R package ape v 5.4 (Paradis et al. 2004).

### Variant calling

We first ran a variant calling analysis, using the *E. lagascae* transcriptome as a reference. We indexed the *E. lagascae* transcriptome using BWA v 0.7.17 (Li and Durbin, 2009) to create a reference and then mapped all the trimmed raw reads to it using the BWA v 0.7.17 “mem” option. We used SAMtools v 1.7 (Li et al., 2009) to convert and sort the alignment files. We then called SNPs using the SAMtools v 1.7 “mpileup” command. Lastly, we used bcftools v 1.9 to filter the SNPs (Narasimhan et al., 2016), running the SAMtools v 1.7 Perl script “vcfutils.pl VarFilter” with default parameters to filter down the candidate variants and to eliminate false positives.

### Discriminant Analysis of Principal Components (DAPC)

We conducted a Discriminant Analysis of Principal Components (DAPC; Jombart et al., 2010) of the SNP data to group the different genotypes avoiding any prior subjective bias using the R package adegenet v 2.1.3 (Jombart and Ahmed, 2011). DAPC is a multivariate method that identifies and describes clusters of genetically related individuals from large datasets, providing a measure of the optimal number of genetic clusters (K) across a range of K values by using the Bayesian Information Criterion (BIC). We set a range of K values from two to seven since K=7 is the number of different species in our dataset. The existence of significant hybridization and introgression would result in K < 7. To identify the optimal number of K, we selected the model with the lowest BIC.

### Phylogenetic inference of introgression

As a first step to detect introgression events between species pairs, we computed phylogenetic species networks. This approach provides a graphical extension of the phylogenetic tree model, representing the gene flow by edges connecting the OTUS that are likely to be linked by introgression. We used the software PhyloNet v 3.6.9 (Than et al., 2008; Wen et al., 2018), which implements a phylogenetic network method based on the frequencies of rooted trees accounting for incomplete lineage sorting (ILS). To generate the input for PhyloNet, we first ultrametricized the trees obtained previously with IQ-Tree v 1.6.1, using the “nnls” method in the “force.ultrametric” function within the R package phytools v 0.6-99 (Revell, 2012). Due to computational limitations, we inferred the species networks using a maximum pseudo-likelihood method (MPL) (Yu and Nakhleh, 2015). We performed the search five times to avoid getting stuck at local optima. We estimated optimal networks among an optimal computational range of 0 to 15 introgression events, determining the most likely network based on Akaike’s Information Criterion (AIC; Bozdogan, 1987) with the generic function for AIC in R package stats v 3.6.1. As AIC may not provide precise values when using pseudo-likelihood phylogenetic networks (Cao et al., 2019), we also estimated the more optimal network by slope heuristic of log-likelihood values. The optimal network was then visualized with Dendroscope v 3.5.10 (Huson and Scornavacca, 2019).

### ABBA-BABA statistic

To assess gene flow between species, we calculated D-statistics, also known as the ABBA-BABA statistic (Durand et al., 2011). To evaluate introgression among the seven species, we used the software Dsuite v 0.1 (Malinsky, 2019), which allows the assessment of gene flow across large datasets and directly from a variant call format (VCF) file. This algorithm computes the D statistic by considering multiple groups of four populations: P1, P2, P3, and O, grouped in asymmetric trees of the form (((P1, P2), P3), O). The site patterns are ordered such that the pattern BBAA refers to P1 and P2 sharing the derived allele (B-derived allele, A-ancestral allele), ABBA to P2 and P3 sharing the derived allele, and BABA to P1 and P3 sharing the derived allele. The ABBA and BABA patterns are expected to occur with equal frequencies, assuming no gene flow (null hypothesis), while a significant deviation from that suggests possible introgression. To assess whether D is significantly different from zero, D-suite uses a standard block-jackknife procedure (Green et al., 2010; Durand et al., 2011), obtaining approximately normally distributed standard errors. As recommended by Malinsky (2019), we used a conservative approach estimating the statistic Dmin, which gives the lowest D-statistic value in a given trio. We used the ruby script “plot_d.rb” to plot into a heatmap the introgression among all the pairs of samples. To complement these analyses, we computed the Fbranch statistic implemented in Dsuite v 0.1 (Mallinsky et al., 2018, Mallinsky et al., 2019). The statistic allows the identification of gene flow events within specific internal branches of a phylogeny. Thus, evaluating the excess sharing of alleles between one species and the descendant or ancestral species, helping to understand when the gene flow happened. We used the whole chloroplast genomes phylogeny from Osuna-Mascaró et al. (2021) in Newick format to establish a reference phylogeny and specify which species could be more accurately treated as sister species (i.e., as P1 and P2) while always using *E. lagascae* as an outgroup.

### Pollen tube growth

The existence of prezygotic barriers can fully impede interspecific hybridization. Therefore, the existence of such barriers may indicate that gene flow across a given set of species is highly unlikely, while the lack of such barriers may indicate plausible hybridization and introgression. To explore the existence of prezygotic barriers, we carried out a preliminary experiment on the growth of pollen tubes on a reduced set of co-occurring species (Table 1). We collected 20 individual plants of each *E. mediohispanicum, E. bastetanum*, and *E. popovii* from natural populations. We grew the plants in a common garden (University of Granada facilities) and moved them into a greenhouse before flowering to exclude pollinators. When the flowers opened, we performed hand-pollination experiments by tipping the anther with a small stick to remove the pollen and placing it on the stigma of a flower from different species previously emasculated (hybrid crosses) or of a flower from the same species but different populations previously emasculated (intra-specific crosses). Moreover, we emasculated some flowers and hand-pollinated them with their own pollen (forced selfing crosses), and some flowers were not manipulated and left for spontaneous self-pollination (spontaneous selfing crosses).

We collected the pistils after 72 hours and preserved them in ethanol at 4ºC until staining of pollen tubes, following the Mori et al. (2006) protocol with minor modifications. In brief, each pistil was cleaned in 70% EtOH for ten minutes and then moved to 50% EtOH, 30% EtOH, and finally distilled water. We softened the samples by placing them into a small petri dish of 8 M NaOH for one hour at room temperature (as recommended by Kearns and Inouye, 1993). Then, we transferred the pistils to distilled water for ten minutes, and afterward, the stigmas were incubated with 0.1 % aniline blue in phosphate buffer (pH 8.3) for two hours. The final slide preparations were examined under a fluorescence microscope with blue light (410 nm) to observe and measure pollen tube development.

## RESULTS

### Ploidy levels

Flow cytometry revealed a wide variation in genome size and, therefore, in DNA ploidy levels across but also within species (Table 2). We found that all samples of *E. fitzii* and *E. nevadense* were diploid. The other species with yellow corollas, *E. mediohispanicum*, also appeared to be predominantly diploid, although the Em71 population deviated from this pattern being tetraploid. The genome size of *E. lasgacae* also corresponded to that of a diploid, while the other purple corolla species showed ploidy levels higher than diploidy (Table 2). Moreover, ploidy levels differed across populations in two of these species. Populations of *E. bastetanum* varied between 4x and 8x, while in *E. popovii*, the range was even greater, from 4x to 10x. In three cases (Ebb12, En05, and En10; Table 2), it was not possible to establish the ploidy level of the samples, and we used those reported in Blanca et al. (1992).

### Transcriptome assembly and orthology inference

The sequencing results and the corresponding summary statistics of the assembled transcriptomes can be found in Osuna-Mascaró et al. (2021). In summary, we obtained between 104K and 382K different Trinity transcripts, producing between 66K and 235K Trinity isogenes. The total assembled bases ranged from 92 Mbp (in the Em21 population of *E. mediohispanicum*) to 319 Mbp (in En10 population of *E. nevadense*). The number of annotated unigenes ranged between 71,606 (*E. nevadense*, En12) and 197,069 (*E. baeticum*, Ebb10); mean value 146,314.35. The highest proportion of annotated unigenes was obtained using BLASTX to search against the SwissProt reference database. Details of the annotated unigenes using different protein databases can be found in Osuna-Mascaró et al. (2021). OrthoFinder assigned 1,519,064 protein gene sequences (96.4% of total) to 92,984 gene families (orthogroups) (Table S2). Among them, 16,941 orthogroups were shared by all species, and their corresponding gene trees were used for further analyses.

### Phylogenetic trees and population clustering

We inferred a coalescence tree using the 16,941 maximum likelihood gene trees obtained with IQ-Tree as input for ASTRAL (Figures 2 and S1). This species tree was nearly fully resolved with high support, having only four nodes with low quartet scores results (posterior probabilities for these nodes: 0.78, 0.77, 0.70, and 0.53; see Figure S1). In this tree, rooted with *E. lagascae*, the 4× population of *E. mediohispanicum* (yellow corollas; Em71) appeared as the first branching OTU. Three clades, although with low supports, were evident. A clade formed by *E. bastetanum* and *E. baeticum*, both species were having purple corollas; another clade including *E. fitzii* (yellow corollas) and the three populations of *E. popovii* (purple corollas); and the last clade including the populations of *E. nevadense* and the 2x populations of *E. mediohispanicum*, both species with yellow corolla. Although there is some species clustering, not all species appear to be monophyletic, supporting a history of hybridization. Moreover, when comparing the species tree with the whole chloroplast genomes phylogeny (Figure 2), we find clear cytonuclear discordances resulting in a significant SH test result (Diff -ln L= 345426.4, p-value < 0.01). This lack of congruence among both phylogenies also supports the hybridization hypothesis.

**Figure 2.**
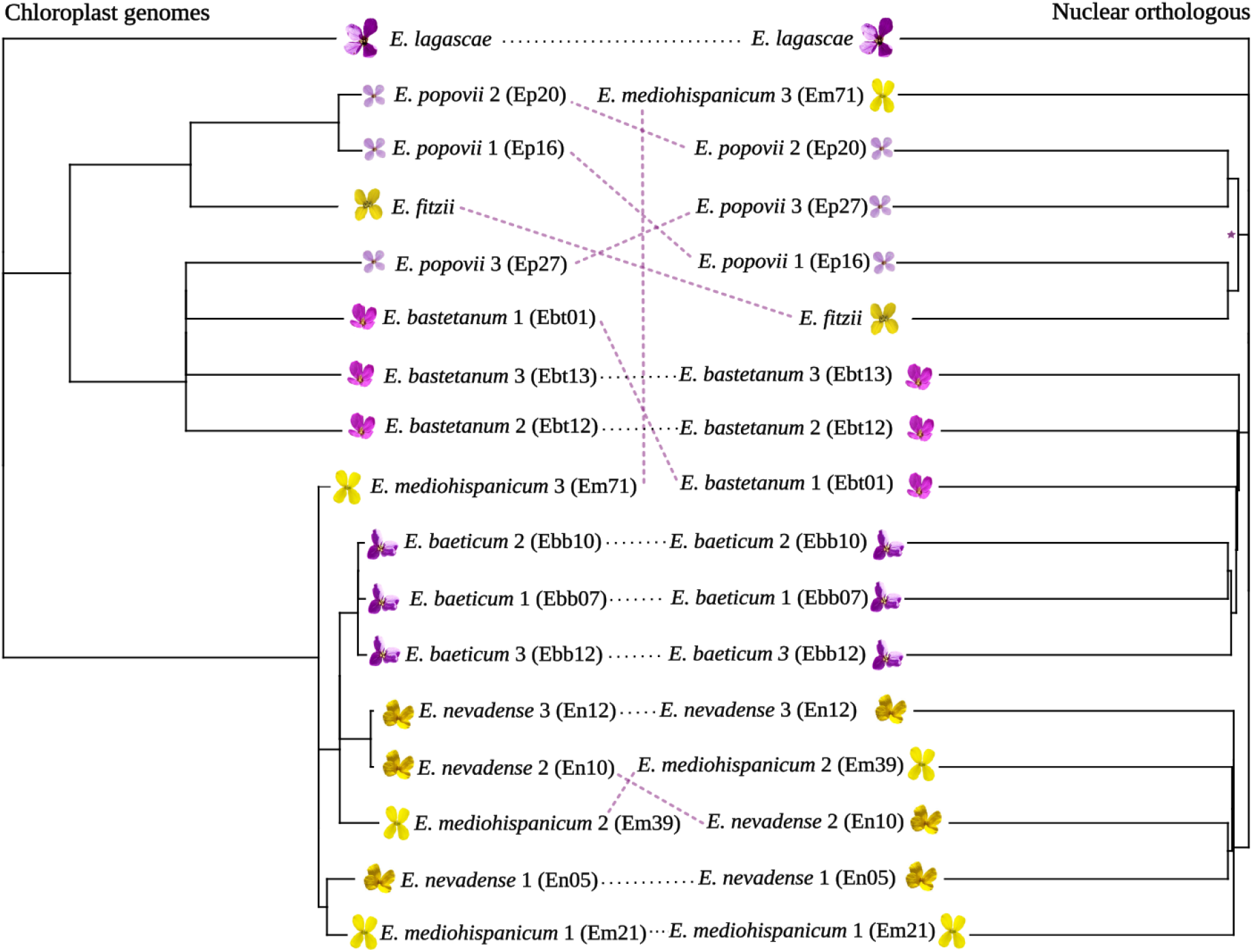
Cyto-nuclear discordance in *Erysimum* spp.. The phylogeny on the left was obtained using whole plastid genomes in Osuna-Mascaró et al. (2021). The phylogeny on the right is a representation of nuclear genome evolution and was generated from the 16,941 maximum likelihood gene trees computed using the SNP data described in the present paper (see text for details).

The discriminant analysis revealed K=4 and K=5 as the most likely number of genetic clusters (Figure 3), both with very similar BIC values (K=4, BIC= 189.99; K=5, BIC= 188.99). The clusters corresponding to K=4 produced the same clusters that appeared in the coalescence tree (Figure 2). However, the clusters corresponding to K=5 included three monospecific groups (for *E. lagascae* -purple corollas-, *E. fitzii* -yellow corollas-, and *E. popovii* -purple corollas-), one for the diploid species with yellow corollas (the three populations of *E. nevadense* and the diploid populations of *E. mediohispanicum*), and the last including all the populations of *E. baeticum* (purple corollas), *E. bastetanum* (purple corollas), *E. popovii* (purple corollas) and Em71, the 4x population of *E. mediohispanicum* (yellow corollas).

**Figure 3.**
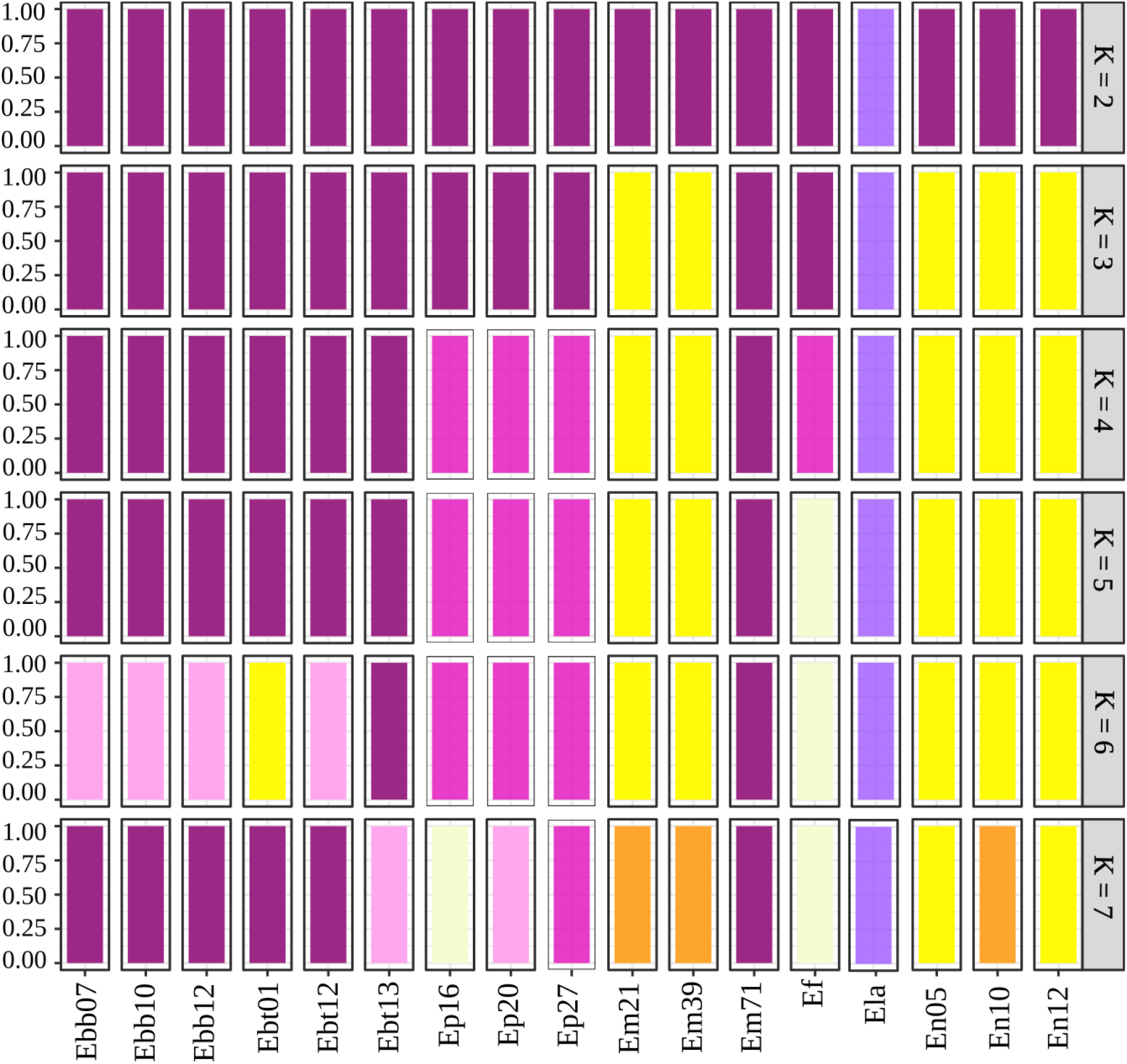
Membership probability plot showing the DAPC results representing the populations grouped into predetermined different clusters (ranging from K=2 to K=7, where each color represents a cluster). The Bayesian analyses (BIC) revealed K = 5 as the most likely number of genetic clusters.

### Analysis of introgression

The network with 13 reticulation instances appeared as the most reliable based on the AIC values for the log-likelihood of the networks (Table S3). The estimates of slope heuristic of log-likelihood values also supported the network with 13 reticulation instances as the most reliable network estimated. This network shows frequent hybridization events in the genealogy of these populations involving yellow and purple species (Figure 4), as indicated by the edges connecting tree branches between different populations and species. Notably, this network includes edges connecting non-terminal branches (see Figure 4), which indicates reticulations with past extinct taxa (i.e., “ghost species”) or incomplete sampled taxa.

**Figure 4.**
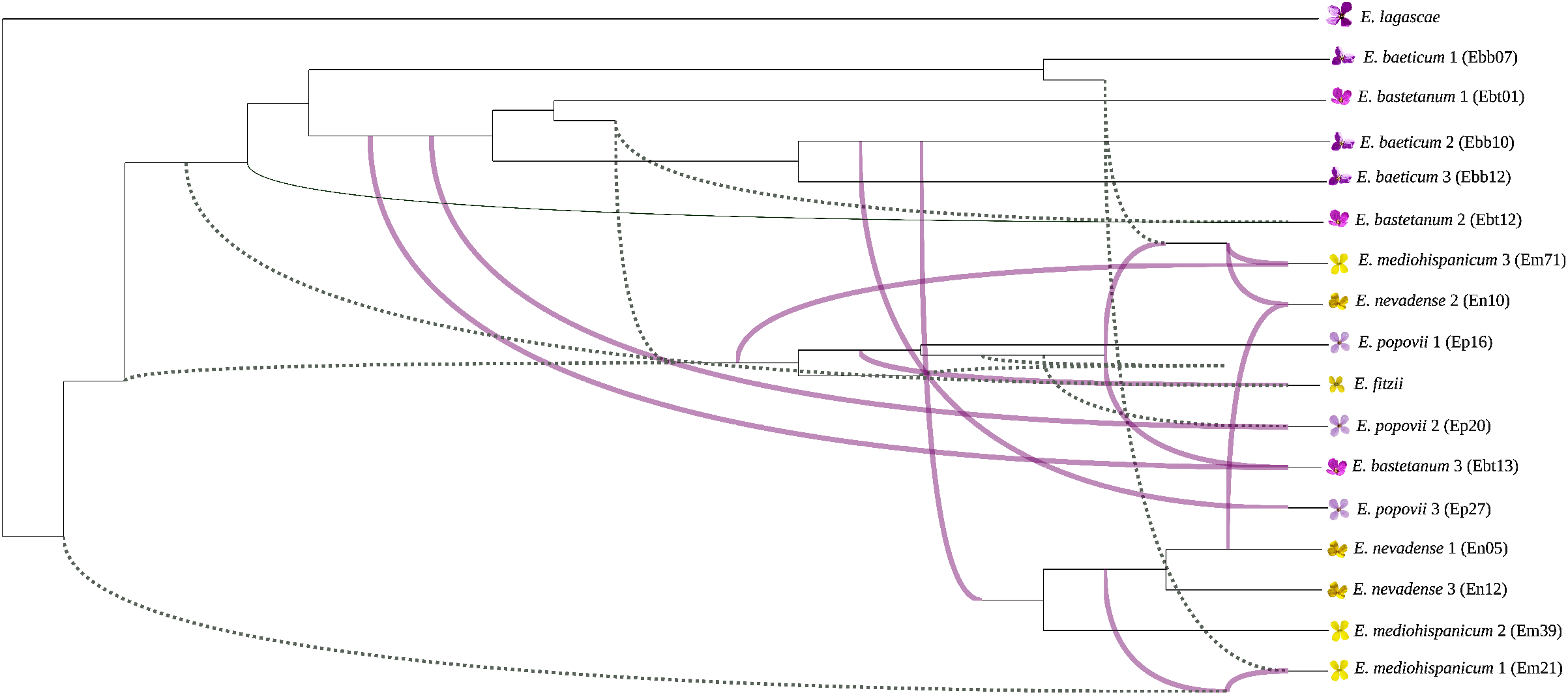
Optimal species network. The graph represents a maximum pseudo-likelihood (MPL) tree with 13 reticulations computed using PhyloNet. These events are represented by edges connecting the tree branches between different individuals and indicate likely hybridization between different taxa. Note that in some instances, introgression appears to involved ancestral or extinct taxa (i.e., ghost species, dotted lines).

The ABBA-BABA analyses support this scenario of frequent hybridization, even using a conservative approach (D-min). We summarized the tested topologies and the inferred D-statistics with corrected p-values for all triplet combinations in Table S4 and Figure S2. The highest signal of introgression occurred between *E. fitzii* (yellow corollas) and *E. baeticum* (purple corollas; populations Ebb12 and Ebb10) and *E. popovii* (purple corollas; Ep16); and between *E. popovii* (purple corollas; Ep16) and *E. bastetanum* (purple corollas; Ebt12) and *E. baeticum* (purple corollas; Ebb07, Ebb10, Ebb12). There was also evidence of interspecific gene flow as manifested by the fbranch statistic (Figure S3) that identifies gene flow events into specific internal branches of the phylogeny while accounting for potential false-positive results due to correlated introgression signatures among closely-related species. Specifically, we found the highest signal of gene flow between *E. bastetanum* (purple corollas; Ebt12) and *E. mediohispanicum* (yellow corollas; Em71, Em21), *E. nevadense* (yellow corollas; En12, En05), and *E. baeticum* (purple corollas; Ebb07, Ebb10); between E. bastetanum (purple corollas; Ebt13) and *E. mediohispanicum* (yellow corollas; Em71) and *E. popovii* (purple corollas; Ep20); between *E. bastetanum* (purple corollas; Ebt01) and *E. mediohispanicum* (yellow corollas; Em21, Em39) and *E. nevadense* (yellow corollas; En12). In addition, we have detected other gene flow events with ancestral or non-sampled taxa (Figure S3).

### Pollen tube growth

A total of 103 preparations of *Erysimum* pistils were examined: 52 from hybrid crosses, 24 from forced selfing crosses, 16 from spontaneous selfing crosses, and 11 from intra-specific crosses (Table S5). Our results showed full growth of pollen tubes (i.e., reaching the ovary) in 63.33 % of intra-specific crosses, 51.92 % of hybrid crosses (χ^2^= 0.50, p-value = 0.513 compared with intra-specific crosses), and only in 29,16 % of forced selfing crosses (χ^2^= 3.73, p-value= 0.074), and in 25,16 % of spontaneous selfing crosses (χ^2^= 4.03, p-value=0.057). Although these last values were not significant, when selfing classes were pooled, it showed a significant reduction in the growth of pollen tubes (χ^2^= 4.93, p-value= 0.039) (Figure S4). Cases in which pollen tubes grew but did not reach the ovary were treated as non-growing. In these cases, we could not estimate whether tube growth had completely stopped or if it was ongoing but developed too slowly to reach the ovary during the duration of the experiment.

## DISCUSSION

Our results suggest that the *Erysimum* species studied here have a strong signature of hybridization and introgression in their genomes. This result is supported by the pollen tube growth experiments that showed that pollen tubes could grow all the way to the ovary in some hybrid crosses, indicating very weak or non-existent prezygotic barriers. Moreover, we found that species with purple flowers are polyploid and have a strong signature of introgression, suggesting an allopolyploid origin. We also found a hybridization signature in the (mostly diploid) yellow species, indicating that hybridization occurred across both colors and ploidy levels.

Several phylogenetic reconstructions have been performed for western Mediterranean *Erysimum* species (e.g., Abdelaziz et al. 2014; Gómez et al. (2014, 2015); Züst et al. (2020)) that have used different strategies (several populations per species or only one representative per species; several nuclear and cytoplasmic sequences or NGS transcriptomic data). Abdelaziz et al. (2014) found that populations of *E. mediohispanicum* and *E. bastetanum*, species analyzed here, did not appear as monophyletic (with some populations placed within other branches of the phylogeny), which was indicative that introgression probably produced important reticulation at the population level. Our analyses support this hypothesis. The reticulate nature of these phylogenies imposes some caution in interpreting phylogenies based on only a few nuclear or cytoplasmic sequences, as suggested by Chan & Levin (2005). In these cases, major divisions may reflect the reality of some old phylogenetic splits. However, it will be challenging for more recent speciation events to obtain a clear picture of the phylogeny without interrogating complete genomes or transcriptomes.

Overall, our results support a hybrid origin for the purple polyploid *Eysimum* Iberian species, as suggested in previous studies (Nieto-Feliner, 1993; Abdelaziz et al., 2014; Gómez et al., 2015b; Osuna-Mascaró, 2020). In particular, we found support for *E. popovii* (purple corollas and polyploid) and *E. fitzii* (yellow corollas and diploid) as sister species. Also, the genome of *E. popovii* exhibited signatures of a hybridization process in which *E. fitzii* may have been involved. The possible hybrid origin of *E. popovii* with *E. fitzii* as a potential parental taxon was previously proposed by Nieto-Feliner (1993) based on morphology. Similarly, a hybrid origin of *E. baeticum* (purple corollas and polyploid) had been previously suggested, in this case with *E. nevadense* (yellow corollas and diploid) implicated as a potential parent (Nieto-Feliner, 1992). Our results showed that these two species appear closely related, and *E. baeticum* may have had an introgression signature of *E. nevadense*. Moreover, our results also suggested a complex scenario for *E. bastetanum* (purple corollas and polyploid), which appears closely related to *E. baeticum* (purple corollas and polyploid). In fact, *E. bastetanum* has been considered a subspecies of *E. baeticum* until recently (Lorite et al., 2014). Therefore, the general pattern that emerged from our results is that these purple species are polyploids of hybrid origin, descending from crosses between an unidentified parent and some diploid, often yellow taxon.

However, our results also suggested a complex evolutionary history for the mostly diploid yellow species. The contributing lineages also often involve unidentified taxa. This might be attributable to insufficient sampling, as we did not include some *Erysimum* species (*E. rondae, E. myriophyllum*, -yellow corollas-, and *E. cazorlense* -purple) that also inhabit the Baetic Mountains, although with a limited distribution (Nieto-Feliner, 1993). At this point, it is impossible to establish whether these taxa may have acted as a source of introgression. In any case, our results show that hybridization and introgression are major contributors to the evolutionary history of this species complex, deserving further research.

Interestingly, we did not find a consistent, predictable pattern of hybridization for most species. Populations of the same species showed differences in their hybridization history, as shown by the ABBA-BABA test (which detected multiple and diverse introgression events) and the PhyloNet reconstructions (which yielded a tree with 13 reticulations as the most optimal network). In the same vein, the DAPC results did not support a scenario with populations clustered by species. Our results are similar to those of previous studies describing asymmetric hybridization patterns as a consequence of differences in ecological pressures across populations and geographical areas (Payton et al., 2019; Sujii et al., 2019; Wang et al., 2019). At this stage, we cannot unambiguously identify any ecological factor behind the asymmetries we detected. However, we did observe variation in pollinators’ preferences and flowering time across populations, which might lead to local differences in gene-flow patterns. Thus, to fully understand this asymmetry in hybridization and why some populations have more introgression signatures than others, future studies considering different ecological pressures for these species and including pollinator censures of wild populations are required.

Evidence of hybridization between at least some of these species has been reported previously (Abdelaziz 2014). Thus, *E. mediohispanicum* and *E. nevadense* show a hybrid zone in a sector of the Spanish Sierra Nevada (Abdelaziz et al., 2021). Pollinators do not appear to constitute strong pre-pollinating barriers since all of these species are extreme generalists and share most pollinators (Gómez et al. 2015b). Moreover, we have found that prezygotic, post-pollination barriers may not be effective since pollen tubes are often growing in hybrid crosses. Contemporary gene flow between different cytotypes of *E. mediohispanicum* seems negligible, as evidenced by an almost complete absence of triploids and other minority cytotypes in the contact zone between tetraploid and diploid populations of this species (Muñoz-Pajares et al. 2018). Historical dynamics of genetic isolation and sympatry might have also played a role (Albaladejo and Aparicio, 2007; Rifkin et al., 2019; Zielinski et al., 2019). These *Erysimum* species are located in a well-known glacial refugium (Médail and Diadema, 2009; Hughes and Woodward, 2017), and thus, the isolation and then re-establishment of gene flow (i.e., secondary contact zones) among populations of different species may have favored locally specific hybridization patterns (Coyne, 2004; Harrison and Larson, 2014; Arnold, 2015). A better knowledge of the historical dynamics of species and populations and past ranges overlap is required to fully understand the genomic pattern of divergence between closely related species. For instance, combining macroecological methods with niche models and phylogenetic approaches could clarify the opportunity for hybridization through evolutionary time (Folk et al., 2018; Aguirre-Liguori et al., 2021).

Furthermore, we detected signatures of ghost introgression, implying that ancestral species have influenced the hybridization history of these *Erysimum* species. This result was first evidenced by cytonuclear discordance, which might be due to past organellar introgression from extinct species (Huang et al., 2014; Folk et al., 2017; Lee-Yaw et al., 2019). We also found a clear signature of ancestral introgression in the phylogenetic species network, in which some of the reticulations appeared from introgression involving “ghost” taxa. Similarly, the fbranch statistic identified gene flow events in internal branches that concurred with introgression with ghost species. Specifically, we observed that some ancestral form of *E. popovii* (purple corollas and polyploid) could have been related to *E. fitzii* (yellow corollas and polyploid). Also, we detected evidence of gene flow between an ancestor of *E. mediohispanicum* (yellow corollas and diploid; Em21), *E. bastetanum* (purple corollas and polyploid), and *E. baeticum* (purple corollas and polyploid). Moreover, the results showed that many past gene flow events could have occurred between *E. baeticum* (purple corollas and polyploid), *E. nevadens*e (yellow corollas and diploid), and *E. bastetanum* (purple corollas and polyploid). In light of these results, it seems that some unidentified ancestral species played a role as introgression sources for both the purple and yellow species. However, as previously noticed, we include only a subset of the Iberian *Erysimum* species in this study; accordingly, we may be mistaking the signal of the unsampled species for that of ancestral taxa. Further research about the ghost introgression’s influence on *Erysimum* evolution, including all the Iberian species and high-quality genome assemblies, would be required to understand the hybridization history thoroughly.

## CONCLUSIONS

Our results indicate that complex evolutionary dynamics have shaped present-day Iberian *Erysimum* diversity. The genomes of extant taxa are the product of multiple polyploidizations, hybridization, and introgression events. Understanding these multi-faceted processes and their interplay is crucial to characterize the evolution of *Erysimum* spp. and probably, of angiosperms in general. Although the evolution of the Iberian *Erysimum* might have been particularly dynamic, this group could be representative of the evolutionary response of multi-species complexes to drastic environmental fluctuations. Further research that incorporates a wider taxonomic sample, whole-genome sequences, and complex demographic and evolutionary statistical methods is needed to precisely characterize the patterns described here.

## Author contributions

COM, RR, JMG, and FP conceived and designed the study. COM analyzed the data with the help of FP, JL, and RH. JL and SC performed the flow cytometry analyses. COM wrote the first draft. The final version of the M.S. was redacted with the contribution of all the authors.

## Acknowledgments

The authors thank Modesto Berbel Cascales, Tatiana López Pérez, Mercedes Sánchez Cabrera, Raquel Sánchez Fernández, Javier Valverde, and Mohamed Abdelaziz, for their help in the lab and fieldwork. Thanks to Pamela Soltis and Douglas Soltis for their help during the first steps of this work. The authors thank the Sierra Nevada National Park headquarters for providing the permits to work in the National Park.

## Funding information

This research is supported by grants from FEDER/Junta de Andalucía-Consejería de Economía y Conocimiento A-RNM-505-UGR18 and P18-FR-3641. This research was also funded by the Spanish Ministry of Science and Innovation (CGL2016-79950-R, CGL2017-86626-C2-2-P), including EU FEDER funds. COM was supported by the Ministry of Economy and Competitiveness (BES-2014-069022).

## List of tables

Table 1. Population code, location, and details of sympatry status for all of the populations sampled.

Table 2. Genome size estimates and ploidy levels of the *Erysimum* species samples included in this study. Genome sizes were estimated with flow cytometry analyses.

## List of figures

Figure 1. Map of the Iberian Peninsula showing the location of the sampled populations.

Figure 2. Cyto-nuclear discordance in *Erysimum* spp.. The phylogeny on the left was obtained using whole plastid genomes in Osuna-Mascaró et al. (2021). The phylogeny on the right is a representation of nuclear genome evolution and was generated from the 16,941 maximum likelihood gene trees computed using the SNP data described in the present paper (see text for details).

Figure 3. Membership probability plot showing the DAPC results representing the populations grouped into predetermined different clusters.

Figure 4. Optimal species network. The graph represents a maximum pseudo-likelihood (MPL) tree with 13 reticulations computed using PhyloNet. These events are represented by edges connecting the tree branches between different individuals and indicate likely hybridization between different taxa. Note that in some instances, introgression appears to involved ancestral or extinct taxa (i.e., ghost species, dotted lines).

## Notes

### Competing Interest Statement

The authors have declared no competing interest.

