## supplementary material for "Hybridization and introgression are prevalent in Southern European *Erysimum* (Brassicaceae) species"

### Supplementary Figures

Figure S1. Phylogeny inferred after a species tree analysis (ASTRAL) of 16,941 gene trees.

Figure S2. Heatmap depicting the introgression found by Dsuite.

Figure S3. Heatmap depicting the gene flow among phylogeny branches estimated with the fbranch statistic.

Figure S4. Pollen tube growth as a result of hand pollination crosses.

### Supplementary Tables

Table S1. Summary of sequencing statistics.

Table S2. Summary of OrthoFinder results.

Table S3. Log likelihood and AIC for all the estimated phylogenetic species network (range of 0 to 15 reticulations).

Table S4. D-statistics with corrected p-values for all triplets combinations (attached as .xls).

Table S5.  *Erysimum* pistils preparations.

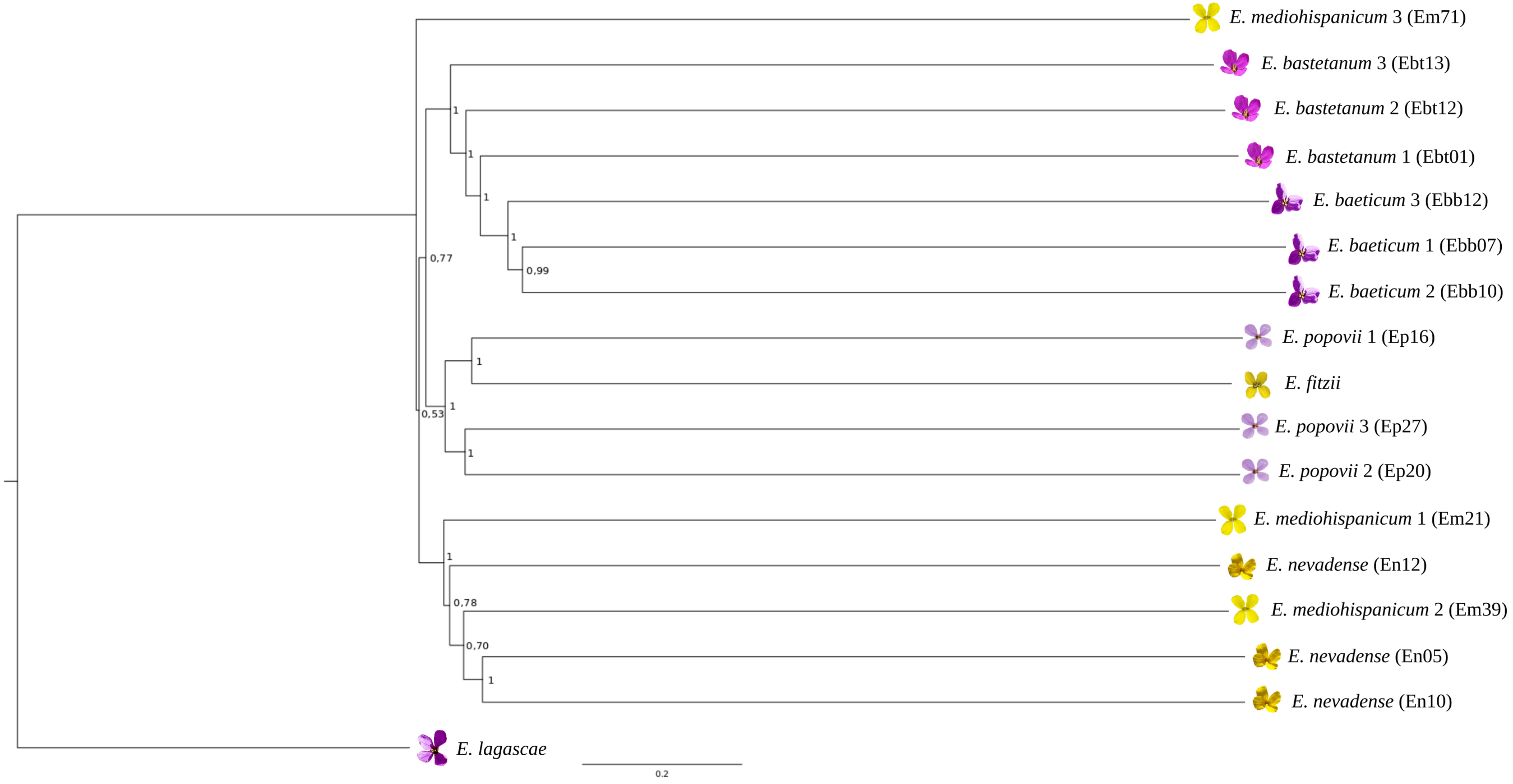


Figure S1. Phylogeny inferred after a species tree analysis (ASTRAL) of 16,941 gene trees. We rooted the obtained species tree to *E. lagascae*, considered the more phylogenetically distant to the rest of species in our data set. Branch support (BS) values are in coalescent units and are a direct measure of the amount of discordance in the gene trees. The code of the populations appears inside parentheses.

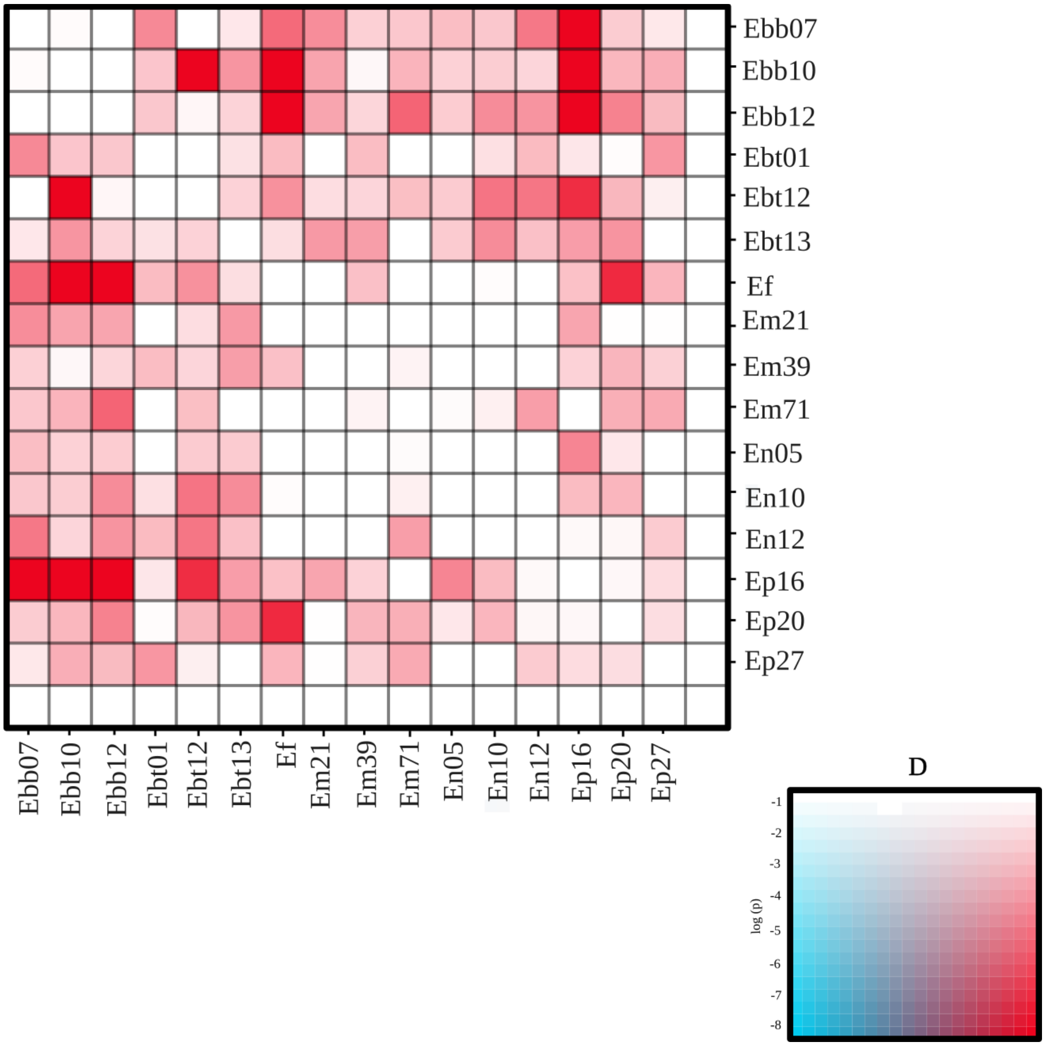


Figure S2. Heatmap depicting the introgression found by Dsuite. The colors of this heatmap show the D-statistic as well as its p-value. Red colors indicate higher D-statistics, and generally more saturated colors indicate greater significance. Thus, the strongest signals for introgression are shown with saturated red, as in the bottom right of the color legend.

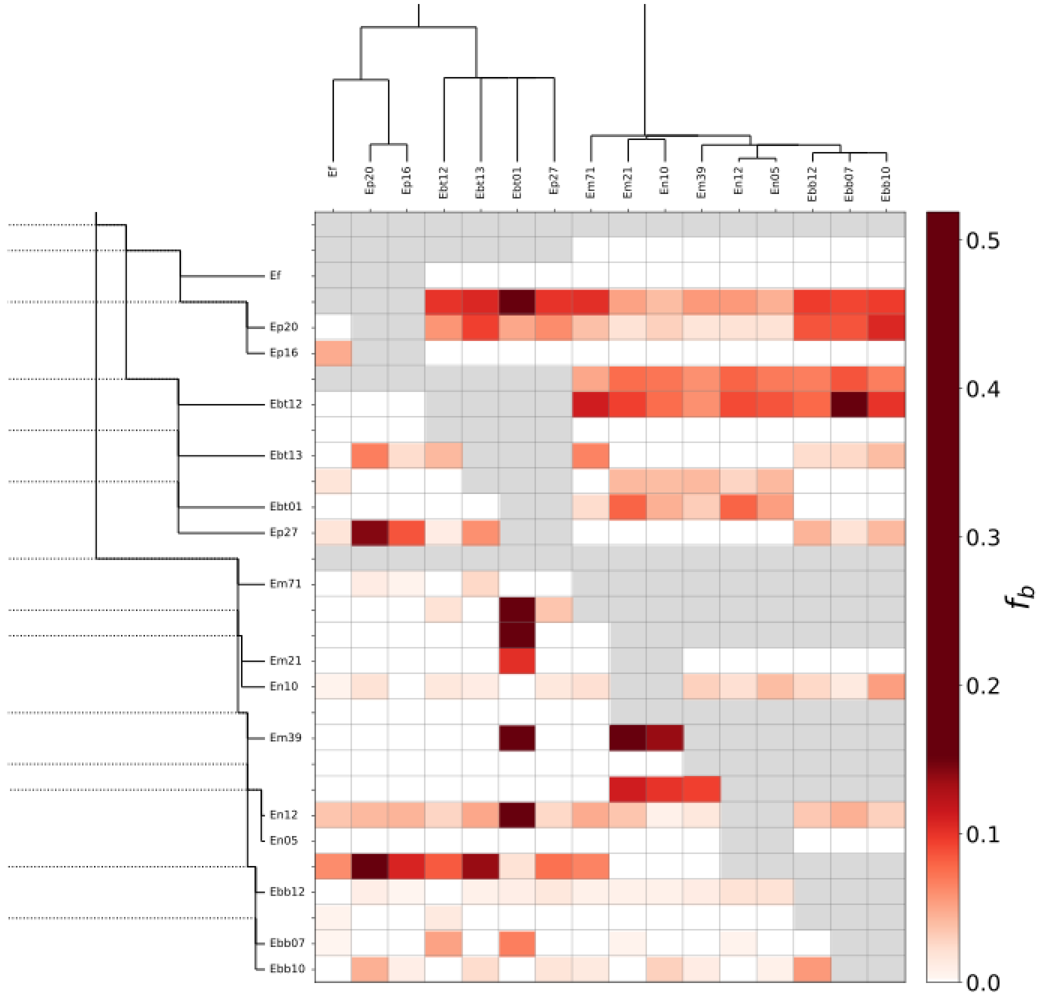


Figure S3. Heatmap depicting the gene flow among phylogeny branches estimated with the fbranch statistic. The phylogenetic tree is shown along the x and y axes (in ‘laddered’ form along the y-axis). The matrix shows the inferred f-branch statistics, showing the gene flow between the branch of the ‘laddered’ tree on the y-axis and the species identified on the x-axis. In the y-axis appears some branches with depicting gene flow that not corresponds to any sampled taxon (horizontal dotted lines). These branches represented ancestral taxa or non-sampled taxa that have contributed to the gene flow.

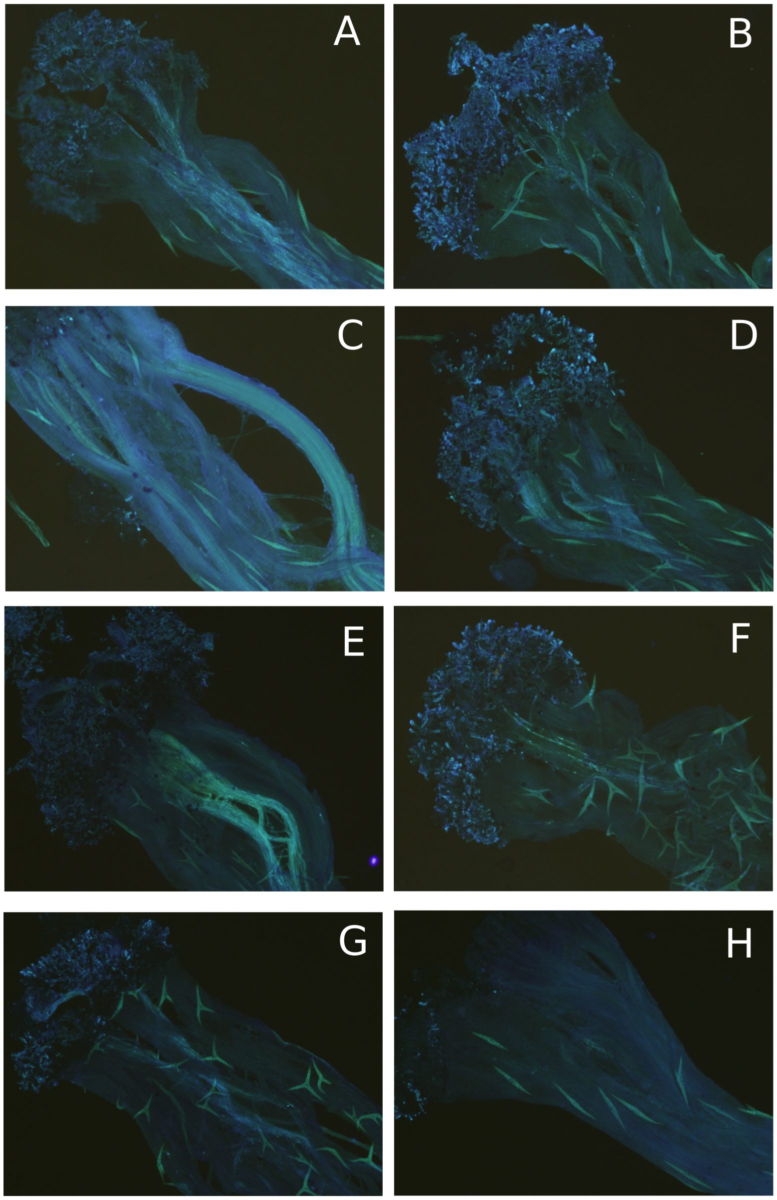

Figure S4. Pollen tube growth as a result of hand pollination crosses. A and B, pollen tube formation after a hybrid cross of *E. bastetanum* with *E. mediohispanicum* (A) and with *E. popovii* (B); C and D, pollen tube formation after a hybrid cross of *E. popovii* with *E. mediohispanicum* (C) and *E. bastetanum* (D); E and F, pollen tube formation after a hybrid cross of *E. mediohispanicum* with *E. popovii* (E) and with *E. bastetanum* (F), F pollen tube formation after an intra-specific cross of *E. popovii* (Ep27) with other population (Ep16) of the same species, and G, shows a lack of growth of the pollen tube.

| Taxon | Population | Total read bases (bp) | Total number of reads (bp) | GC (%) | Q20 (%) | Q30 (%) | Gb | Reads after trimming (bp) |
|---|---|---|---|---|---|---|---|---|
| *E. baeticum* | Ebb07 | 10,342,234,016 | 68,491,616 | 47.38 | 96.13 | 91.88 | 5.1 | 60,985,000 |
| *E. baeticum* | Ebb10 | 20,383,113,406 | 134,987,506 | 47.25 | 96.63 | 92.11 | 9.9 | 119,520,613 |
| *E. baeticum* | Ebb12 | 20,634,741,014 | 136,653,914 | 47.30 | 96.71 | 92.24 | 10.1 | 121,103,833 |
| *E. bastetanum* | Ebt01 | 11,454,840,974 | 75,859,874 | 44.88 | 99.89 | 99.10 | 5.0 | 75,854,979 |
| *E. bastetanum* | Ebt12 | 10,417,333,262 | 68,988,962 | 45.71 | 98.22 | 95.24 | 5.2 | 65,757,353 |
| *E. bastetanum* | Ebt13 | 10,777,576,680 | 71,374,680 | 45.47 | 98.58 | 95.91 | 5.4 | 65,765,324 |
| *E. fitzii* | Ef01 | 10,156,440,596 | 67,261,196 | 47.09 | 98.94 | 96.66 | 4.6 | 66,245,223 |
| *E. lagascae* | Ela07 | 7,201,775,508 | 71,304,708 | 43.50 | 96.47 | 93.68 | 5.4 | 65,740,933 |
| *E. mediohispanicum* | Em21 | 10,235,928,808 | 67,787,608 | 48.86 | 96.05 | 91.72 | 4.7 | 60,130,101 |
| *E. mediohispanicum* | Em39 | 23,224,849,148 | 153,806,948 | 47.49 | 96.79 | 92.43 | 11.2 | 136,858,973 |
| *E. mediohispanicum* | Em71 | 10,242,859,708 | 67,833,508 | 46.51 | 98.18 | 95.14 | 5.0 | 64,619,740 |
| *E. nevadense* | En05 | 22,265,032,144 | 147,450,544 | 46.73 | 96.65 | 92.20 | 10.8 | 130,509,040 |
| *E. nevadense* | En10 | 24,045,140,340 | 159,239,340 | 44.94 | 96.90 | 92.72 | 11.4 | 142,305,123 |
| *E. nevadense* | En12 | 10,141,853,090 | 67,164,590 | 47.49 | 96.17 | 91.96 | 5.1 | 59,838,058 |
| *E. popovii* | Ep16 | 10,556,586,670 | 69,911,170 | 46.26 | 96.09 | 91.78 | 5.7 | 62,135,880 |
| *E. popovii* | Ep20 | 10,860,711,844 | 71,925,244 | 45.98 | 98.12 | 94.96 | 5.5 | 68,273,570 |
| *E. popovii* | Ep27 | 25,555,407,006 | 169,241,106 | 47.38 | 96.64 | 92.14 | 12.5 | 149,733,87 |

Table S1. Summary of the sequencing statistics.

| | |
|---|---|
| Total number of Orthogroup | 92,984 |
| Mean Orthogroup size | 16.3 |
| Median Orthogroup size | 7.0 |
| Number of protein genes | 1,574,983 |
| N. of genes in Orthogroups | 1,519,064 |
| N. of unassigned protein genes | 55,919 |
| Percentage of genes in Orthogroups | 96.4 |
| Percentage of unassigned protein genes | 3.6 |
| Number of Orthogroup with all species present | 16,941 |

Table S2. Summary of OrthoFinder results.

| | Log likelihood | AIC |
|---|---|---|
| 1 | -1.0224071648028404E7 | 20448145 |
| 2 | -1.022316938865218E7 | 20446343 |
| 3 | -1.0222191339585347E7 | 20444389 |
| 4 | -1.0221615102393162E7 | 20443238 |
| 5 | -1.022131616996915E7 | 20442642 |
| 6 | -1.0220690757836848E7 | 20441394 |
| 7 | -1.0220980513494018E7 | 20441975 |
| 8 | -1.0223136397700764E7 | 20446289 |
| 9 | -1.0220246575984979E7 | 20440511 |
| 10 | -1.0220445144143904E7 | 20440910 |
| 11 | -1.0220701238338212E7 | 20441414 |
| 12 | -1.0219898642573046E7 | 20439821 |
| 13 | -1.0219513355204735E7 | *20439053 |
| 14 | -1.021989388332834E7 | 20439816 |
| 15 | -1.0220243030447358E7 | 20440516 |

Table S3. Log likelihood and AIC for all the estimated phylogenetic species network (range of 0 to 15 reticulations). The network with the lower AIC is depicted by an asterisk.

| Hybrid crosses | Pollen tube full growth |
|---|---|
| *E. mediohispanicum* x *E. bastetanum* | |
| Em08 x Ebt13 | Yes |
| Em08 x Ebt13 | Yes |
| Em08 x Ebt13 | No |
| Em21 x Ebt01 | No |
| Em21 x Ebt13 | Yes |
| Em71 x Ebt13 | No |
| *E. bastetanum* x *E. mediohispanicum* | |
| Ebt01 x Em08 | Yes |
| Ebt01 x Em21 | Yes |
| Ebt01 x Em21 | Yes |
| Ebt01 x Em21 | Yes |
| Ebt01 x Em21 | No |
| Ebt13 x Em08 | Yes |
| Ebt13 x Em08 | Yes |
| Ebt13 x Em21 | No |
| Ebt13 x Em71 | No |
| *E. mediohispanicum* x *E. popovii* | |
| Em08 x Ep14 | Yes |
| Em08 x Ep14 | No |
| Em08 x Ep27 | Yes |
| Em08 x Ep27 | No |
| Em08 x Ep27 | No |
| Em21 x Ep14 | Yes |
| Em21 x Ep14 | No |
| Em21 x Ep14 | No |
| Em21 x Ep14 | No |
| Em21 x Ep27 | Yes |
| Em71 x Ep27 | Yes |
| Em71 x Ep27 | No |
| Em71 x Ep27 | No |
| Em71 x Ep27 | No |
| *E. popovii x E. mediohispanicum* | |
| Ep14 x Em08 | Yes |
| Ep14 x Em21 | No |
| Ep14 x Em21 | No |
| Ep27 x Em21 | Yes |

| | |
|---|---|
| Ep27 x Em21 | Yes |
| Ep27 x Em21 | No |
| Ep27 x Em21 | No |
| Ep27 x Em08 | Yes |
| Ep27 x Em08 | Yes |
| Ep27 x Em08 | Yes |
| Ep27 x Em08 | Yes |
| Ep27 x Em08 | No |
| Ep27 x Em08 | No |
| *E. popovii x E. bastetanum* | |
| Ep14 x Ebt01 | No |
| Ep27 x Ebt01 | No |
| Ep27 x Ebt13 | No |
| *E. bastetanum x E. popovii* | |
| Ebt01 x Ep14 | Yes |
| Ebt13 x Ep14 | Yes |
| Ebt13 x Ep27 | Yes |
| Ebt13 x Ep27 | Yes |
| Ebt13 x Ep27 | No |
| Ebt01 x Ep27 | Yes |
| Ebt13 x Ep27 | Yes |
| Forced selfing | |
| Em08 | No |
| Em21 | Yes |
| Em21 | Yes |
| Em21 | No |
| Em21 | No |
| Em21 | No |
| Em21 | No |
| Em21 | No |
| Em21 | No |
| Em71 | No |
| Ebt01 | No |
| Ebt13 | Yes |
| Ebt13 | No |
| Ebt13 | No |
| Ep27 | Yes |
| Ep27 | Yes |
| Ep14 | Yes |
| Ep27 | No |

| | |
|---|---|
| Ep27 | No |
| Ep14 | No |
| Ep27 | Yes |
| Ep27 | No |
| Ep27 | No |
| Ep27 | No |
| Spontaneous selfing crosses | |
| Em08 | Yes |
| Em21 | Yes |
| Em21 | No |
| Em21 | No |
| Em21 | No |
| Em21 | No |
| Em71 | Yes |
| Ebt01 | No |
| Ebt01 | No |
| Ebt13 | No |
| Ep27 | No |
| Ep27 | No |
| Ep27 | Yes |
| Ep27 | No |
| Ep27 | No |
| Ep27 | No |
| Intra-specific crosses | |
| Em08 x Em21 | Yes |
| Em08 x Em21 | Yes |
| Em21 x Em08 | Yes |
| Em21 x Em21 | Yes |
| Em21 x Em71 | Yes |
| Em71 x Em08 | Yes |
| Em71 x Em08 | No |
| Ebt13 x Ebt13 | No |
| Ebt13 x Ebt13 | No |
| Ep27 x Ep27 | Yes |
| Ep27 x Ep27 | No |

Table S5. A total of 103 *Erysimum* pistils preparations were examined: 52 from hybrid crosses, 24 from forced selfing crosses, 16 from spontaneous selfing crosses, and 11 from intra-specific crosses. *E. bastetanum* (Ebt01, Ebt13), *E. mediohispanicum* (Em08, Em21, Em71), *E. popovii* (Ep14, Ep27)
